# geneSync: Gene Symbol Harmonization for Large-scale RNA-seq Data Integration

**DOI:** 10.64898/2026.05.04.722831

**Authors:** Zhijun Feng, Ting Li

**Author notes:** Correspondence: Ting Li, E-mial.

## Abstract

Cross-cohort integration of transcriptomic data is a routine strategy for boosting statistical power and enhancing generalizability. However, gene nomenclature inconsistencies across datasets—arising from annotation version updates, historical renaming, and synonym reassignment—introduce silent mismatches during feature alignment, causing genes to be falsely classified as absent or split into duplicate features. Here, we present geneSync, an R package that performs gene symbol harmonization as a quality-control (QC) step prior to data integration. geneSync uses a hierarchical matching strategy, prioritizing exact matches to authoritative gene symbols, then exact matches to National Center for Biotechnology Information (NCBI) gene symbols, and finally synonym-based fallback. It includes built-in offline databases for human, mouse, and rat, and supports auditable conflict resolution, cross-species ortholog mapping, and native integration with Seurat and SingleCellExperiment objects. Benchmarking across six mouse hippocampus scRNA-seq datasets spanning 2020–2025 and five CellRanger versions shows that 1.41%–6.22% of features require synonym resolution, and harmonization improves pairwise gene overlap by up to 13.14 percentage points, rescuing 707–1,098 genes per dataset pair. Notably, CellRanger annotation version—rather than data collection year—was identified as the primary driver of nomenclature discrepancy. geneSync is freely available at https://github.com/xiaoqqjun/geneSync.

## Introduction

RNA sequencing and its derivative technologies—bulk RNA-seq, single-cell RNA-seq (scRNA-seq), single-nucleus RNA-seq (snRNA-seq), and spatial transcriptomics—have enabled systematic investigation of tissue architecture and disease heterogeneity [1]. As public transcriptomic data continue to expand, cross-study data integration has become a standard strategy for improving statistical power and supporting cross-cohort validation [2].

The reliability of integrative analyses depends on a fundamental prerequisite: datasets must share a consistent feature coordinate system. In practice, gene nomenclature is not static. Reference genomes and annotation frameworks undergo iterative updates, genes are officially renamed, synonyms are reassigned, and capitalization or punctuation conventions are revised [3]. Different studies frequently employ different versions of GENCODE, Ensembl, or RefSeq annotations, and critically, different versions of processing pipelines such as CellRanger ship with different Gene Transfer Format (GTF) annotation files [4]. Consequently, the same biological gene may appear under different symbols across datasets.

This inconsistency produces three types of systematic consequences during feature alignment: (1) genuinely present genes are misclassified as absent, shrinking gene intersections; (2) the same gene appears under different symbols, generating duplicate features that alter count distributions; and (3) multiple historical symbols are indiscriminately collapsed onto the same recommended symbol, introducing untraceable bias. These errors seldom trigger explicit warnings, yet propagate into differential expression, pathway enrichment, cell-type annotation, and integrative embedding [5,6].

While re-aligning raw data to a unified reference genome is the most rigorous approach, raw FASTQ files are often unavailable, and uniform re-alignment imposes substantial computational demands. Existing gene identifier conversion tools—including biomaRt [7], org.Hs(Mm).eg.db [8,9], and HGNChelper [10], address parts of this problem but have notable limitations for integration-oriented quality control (QC): biomaRt requires network access and is subject to API instability; org.Hs.eg.db does not support synonym matching; and none provide integrated conflict logging or direct object-level deployment for single-cell workflows (Table 1).

**Table 1.**
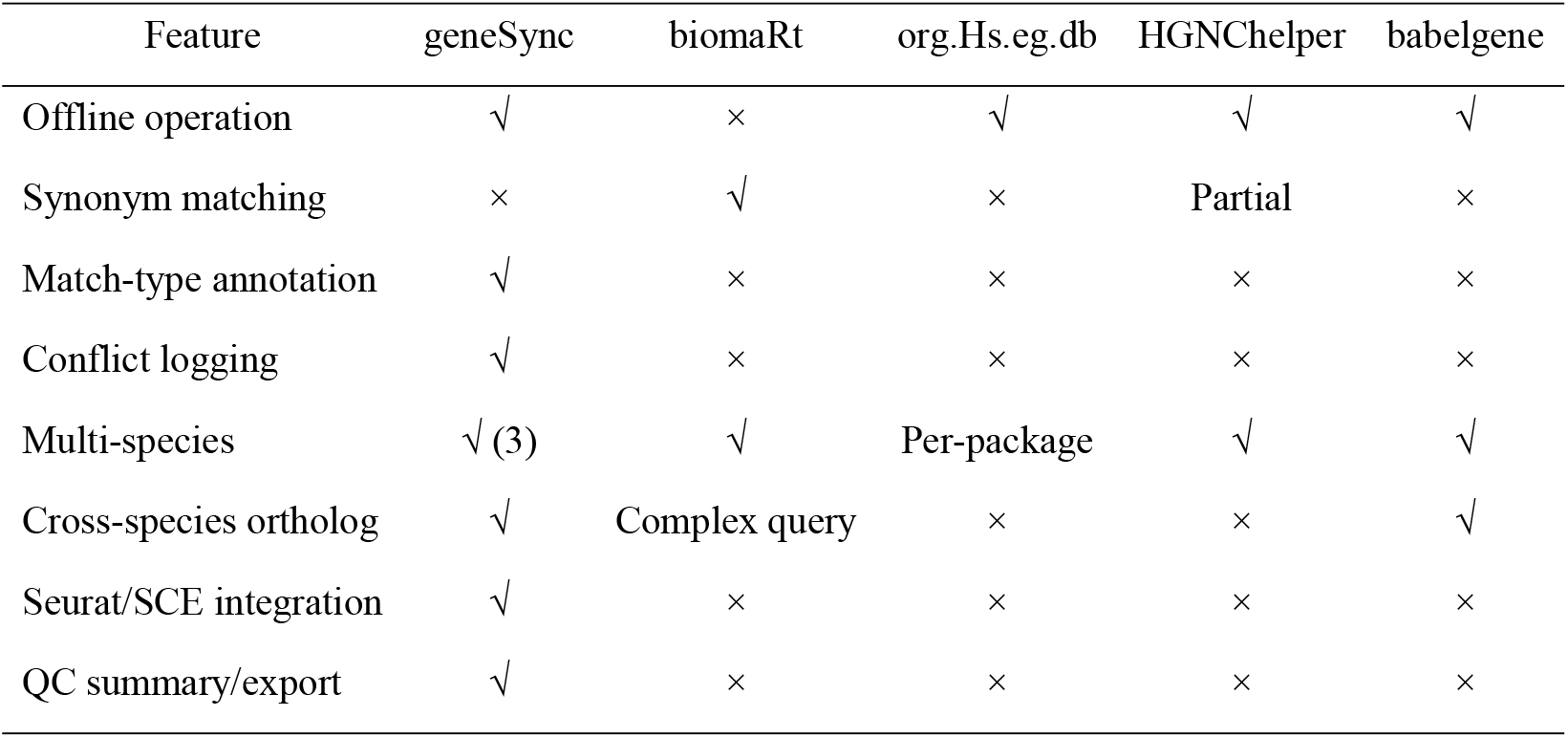
Feature comparison of geneSync with existing gene annotation tools.

| Feature | geneSync | biomaRt | org.Hs.eg.db | HGNChelper | babelgene |
| --- | --- | --- | --- | --- | --- |
| Offline operation | √ | × | √ | √ | √ |
| Synonym matching | × | √ | × | Partial | × |
| Match-type annotation | √ | × | × | × | × |
| Conflict logging | √ | × | × | × | × |
| Multi-species | √ (3) | √ | Per-package | √ | √ |
| Cross-species ortholog | √ | Complex query | × | × | √ |
| Seurat/SCE integration | √ | × | × | × | × |
| QC summary/export | √ | × | × | × | × |

Here, we present geneSync, an R package that performs gene symbol harmonization as a prerequisite QC step for RNA-seq data integration. geneSync provides offline hierarchical matching, auditable conflict resolution, cross-species ortholog mapping, and native integration with Seurat and SingleCellExperiment objects. We demonstrate through benchmarking on six public scRNA-seq datasets spanning five years and five CellRanger versions that symbol inconsistency is prevalent and consequential, and that harmonization substantially improves feature overlap.

## Methods

### Offline mapping databases

geneSync relies on built-in offline mapping tables constructed from versioned snapshots of the NCBI Gene database (https://www.ncbi.nlm.nih.gov/gene; snapshot date: February 5, 2026) [11]. Three species are supported: human (*Homo sapiens*; 193,859 records), mouse (*Mus musculus*; 113,674 records), and rat (*Rattus norvegicus*; 66,389 records). Each record contains seven fields: Entrez Gene ID, NCBI symbol, authoritative nomenclature symbol (HGNC/MGI/RGD), Ensembl ID, chromosomal location, synonyms (pipe-delimited historical aliases), and gene type. Cross-species mapping is based on the NCBI gene_orthologs database, providing one-to-one ortholog pairs for human–mouse (16,822 pairs), human–rat (16,753 pairs), and mouse–rat (16,494 pairs, bridged through human).

### Hierarchical matching strategy

The core function gene_convert() maps input symbols to standardized identifiers through a three-tier hierarchical strategy. **Tier 1 (exact authoritative symbol match):** Input symbols are matched against species-specific nomenclature authority symbols (HGNC/MGI/RGD) using a pre-built lookup table; matching is case-insensitive by default. **Tier 2 (exact NCBI symbol match):** Unresolved inputs from Tier 1 are matched against the NCBI symbol field, with the corresponding authoritative symbol backfilled when available. **Tier 3 (synonym fallback):** Only inputs unresolved by both preceding tiers are searched against the synonym field via regular expression matching; matches are explicitly flagged as synonym in the output. For each input, the function returns a standardized result (final_symbol) following the priority: authoritative symbol > NCBI symbol > original input (when unmatched), together with the match type (exact_authority, exact_symbol, synonym, or not_found) and all traceable fields including Entrez Gene ID, Ensembl ID, and chromosomal location.

### Many-to-one conflict resolution

When multiple input symbols map to the same final_symbol—for example, due to a RIKEN identifier and its official symbol coexisting in the feature list—duplicate features arise that must be resolved before downstream analysis. geneSync adopts different strategies for bulk and single-cell workflows to match their distinct analytical contexts.

For bulk RNA-seq data, gene_convert() returns a standardized mapping table that users merge with their original expression matrix. Because bulk analyses typically operate on individual datasets rather than multi-platform integration, and because the optimal duplicate-handling strategy depends on the specific analytical context (e.g., summing counts vs. retaining the higher-expressed copy), geneSync does not impose automatic collapsing at this stage. Instead, duplicate entries are explicitly flagged in the output table, enabling users to apply context-appropriate resolution.

For single-cell data, where multi-dataset integration is routine and feature lists must be unique within each Seurat or SingleCellExperiment object, geneSync provides automated resolution within gene_sync_add_to_obj(). When multiple input features map to the same final_symbol, the function evaluates two metrics—the proportion of cells expressing the gene (detection rate) and mean expression level—to retain the most informative feature. Remaining duplicates are flagged as keep = “no” with their resolution rationale recorded. All decisions are stored in a traceable conversion table exportable via export_gene_sync_table(), ensuring that no collapsing occurs silently.

### Cross-species ortholog mapping

The function gene_ortholog() converts gene symbols between species using pre-built one-to-one ortholog lookup tables. Input symbols are first resolved to Entrez Gene IDs via gene_convert(), then mapped to the target species through vectorized hash-table lookup (O(n) complexity), achieving approximately 1–2 seconds for 18,000 genes compared to 50–70 seconds in naive loop implementations. A batch wrapper batch_ortholog() provides summary statistics for large-scale conversions.

### Integration with expression objects

geneSync provides the function gene_sync_add_to_obj() for direct integration with Seurat objects (v4/v5) and SingleCellExperiment objects [12,13]. This function performs symbol harmonization, duplicate collapsing, and object consistency maintenance in a single call. The complete conversion table and parameters are stored in retrievable object slots (misc for Seurat; metadata for SingleCellExperiment), ensuring full traceability. Summary reporting is available via print_gene_sync_summary() and convert_summary(), outputting match rates, conflict counts, and timestamped logs.

Additionally, geneSync maintains equivalence relationships for mitochondrial gene nomenclature (e.g., Atp6/mt-Atp6 for mouse; ATP6/MT-ATP6 for human), ensuring correct detection by pattern-based QC filters when input features use non-prefixed NCBI symbols. Auxiliary functions include get_mito_genes() for retrieving species-specific mitochondrial gene lists and get_chr_genes() for chromosome-specific gene queries.

geneSync is implemented in R (≥3.5.0) with dependencies on Seurat, Matrix, and dplyr, and is available at “https://github.com/xiaoqqjun/geneSync“ under the MIT license.

### Benchmark datasets

To evaluate the prevalence and impact of gene symbol inconsistency, we assembled six mouse hippocampus scRNA-seq datasets spanning five years (2020–2025) and five CellRanger versions (Table 2). Four datasets were obtained from Gene Expression Omnibus (GEO) database (GSE157985 [14], GSE212576 [15], GSE235490 [16], GSE270158 [17]) and two were generated in-house using CellRanger V7.0 and V10.0. This design enables assessment of nomenclature drift driven by both annotation version updates and CellRanger pipeline evolution.

**Table 2.** Benchmark datasets and symbol harmonization results.

| GEO ID | Year | CellRanger | Cells | Raw features | Exact (%) | Synonym (%) | Not found (%) | Dup.symbols |
| --- | --- | --- | --- | --- | --- | --- | --- | --- |
| GSE157985 | 2020 | V3.0 | 13,275 | 23,052 | 85.64 | 4.78 | 9.58 | 30 |
| GSE212576 | 2022 | V4.0 | 47,583 | 25,417 | 85.87 | 4.04 | 10.09 | 35 |
| GSE235490 | 2023 | V1.1 | 88,623 | 25,488 | 85.55 | 4.58 | 9.88 | 34 |
| GSE270158 | 2024 | V3.0 | 15,241 | 19,055 | 88.90 | 6.22 | 4.88 | 33 |
| Own_V7 | 2025 | V7.0 | 9,667 | 21,770 | 88.44 | 3.41 | 8.15 | 27 |
| Own_V10 | 2025 | V10.0 | 13,000 | 25,796 | 86.52 | 1.41 | 12.07 | 28 |

## Results

### Prevalence of nomenclature inconsistency across CellRanger versions

We applied geneSync to each of the six datasets independently (Table 2, Table S1). The proportion of features resolved through exact authoritative symbol matching ranged from 85.55% to 88.90%, while 1.41%–6.22% of features required synonym resolution to be mapped to current recommended symbols. The remaining 4.88%–12.07% of features could not be matched, representing non-coding RNAs, predicted genes, or platform-specific identifiers absent from the NCBI Gene database.

Notably, synonym match rates varied more strongly with CellRanger version than with collection year. The dataset processed with CellRanger V10.0 (2025) exhibited the lowest synonym rate (1.41%, 363 genes), whereas CellRanger V3.0 (2024) showed the highest (6.22%, 1,185 genes). Two datasets collected in the same year (2025) using different CellRanger versions (V7.0 and V10.0) differed by 2.0 percentage points in synonym rate (3.41% vs. 1.41%). This pattern indicates that the GTF annotation files bundled with different CellRanger versions are the primary driver of symbol inconsistency, rather than the year of data collection per se. Even in the most recent dataset (CellRanger V10.0), 363 features still required synonym resolution, confirming that nomenclature drift does not resolve spontaneously with pipeline updates.

Each dataset contained 27–35 genes where multiple input symbols mapped to the same recommended symbol (duplicate symbols). Across all six datasets, 188 unique duplication events were identified (Table S2), predominantly involving RIKEN clone identifiers and predicted gene symbols being reclassified to official nomenclature (e.g., Gm28286 → A830008E24Rik; 3110035E14Rik → Vxn; Kdelc1 → Poglut2). This pattern reflects the ongoing reclassification of provisional gene symbols in the mouse genome, which constitutes the primary source of nomenclature drift across annotation versions.

### Impact on feature overlap in multi-dataset integration

To assess practical consequences, we computed pairwise feature overlap for all 15 dataset pairs before and after harmonization (Table S3). Before harmonization, pairwise gene overlap (Jaccard index) ranged from 61.18% to 92.73%. After applying geneSync, overlap improved to 73.98%–93.46%, representing gains of 0.73–13.14 percentage points. Per comparison, 707–1,098 genes were rescued— genes present in both datasets but recorded under different symbols.

The magnitude of improvement showed a clear association with CellRanger version discrepancy rather than year gap alone. The largest improvement (13.14 percentage points, 1,069 genes rescued) occurred between CellRanger V3.0 (2024) and V10.0 (2025)—datasets separated by only one year but employing substantially different annotation files. Conversely, datasets processed with similar CellRanger versions showed minimal improvement regardless of year gap (e.g., CellRanger V4.0 vs. V1.1: 0.73 percentage points). This finding underscores that annotation pipeline version is a stronger predictor of nomenclature discrepancy than data vintage.

For the six-way intersection across all datasets, the common feature set expanded from 15,763 genes (raw) to 16,151 genes (harmonized), with 962 genes rescued into the shared feature space (Table S4). The intersection-to-union ratio improved from 51.51% to 64.93%. In large-scale integration where multiple cohorts are combined, these per-pair losses compound: genes inconsistently named in even one dataset are excluded from the intersection used for integration anchors, highly variable gene selection, and joint embedding.

Among the rescued genes, the majority were RIKEN identifiers and predicted gene symbols that had been reclassified to official MGI nomenclature between annotation versions (e.g., Fam126b → Hycc2; Als2cr12 → Flacc1; Sgol2a → Sgo2a) (Table S5-S6). While these genes are not typically considered as canonical cell-type markers, they include protein-coding genes that contribute to expression variance calculations, dimensionality reduction, and integration anchor selection, and their systematic exclusion introduces non-random bias into the feature space.

### Comparison with existing tools

We compared geneSync with commonly used gene annotation tools across key capability dimensions (Table 1). geneSync uniquely combines offline operation, synonym support with explicit match-type annotation, many-to-one conflict logging, cross-species ortholog mapping, and direct single-cell object integration. For a typical input of 20,000 genes, gene_convert() completes in approximately 5-10 seconds; gene_ortholog() processes 18,000 genes in 10-20 seconds. Runtime scales approximately linearly with input feature count.

## Discussion

Our benchmarking across six mouse hippocampus scRNA-seq datasets demonstrates that gene symbol inconsistency is prevalent and introduces systematic bias at the feature alignment stage. Synonym resolution rates of 1.41%–6.22%, corresponding to 363–1,185 genes per dataset, translate into measurable losses in pairwise gene overlap (up to 13.14 percentage points) and the introduction of 27– 35 duplicate features per dataset. In a six-way integration, 962 genes were rescued into the shared feature space after harmonization.

A key finding of this study is that CellRanger version—rather than data collection year—is the primary driver of nomenclature discrepancy. Different CellRanger releases bundle different GTF annotation files derived from different Ensembl/GENCODE versions, resulting in systematic differences in gene symbol assignments. This observation has practical implications: researchers integrating datasets processed with different pipeline versions should perform symbol harmonization regardless of how recently the data were generated.

The predominance of RIKEN clone identifiers and predicted gene symbols (e.g., Gm* and *Rik) among synonym-resolved features reflects the ongoing reclassification of provisional gene symbols in the mouse genome by the Mouse Genome Informatics (MGI) consortium [18]. While these genes are rarely used as canonical cell-type markers, their systematic exclusion from integration feature sets introduces non-random bias into variance calculations, anchor selection, and joint embedding. geneSync addresses this by providing explicit match-type classification and auditable conflict resolution, transforming what would otherwise be silent data loss into a documented QC decision.

We note that in datasets processed by standard pipelines such as CellRanger, mitochondrial genes are already annotated with species-appropriate prefixes (MT-for human, mt-for mouse), and thus typically do not require symbol conversion for QC metric calculations. The mitochondrial gene handling in geneSync is primarily relevant when working with gene lists derived from direct database queries, non-standard annotation sources, or legacy datasets processed with custom pipelines.

Several additional limitations should be acknowledged. First, the current offline databases cover human, mouse, and rat; users working with other species will need custom mapping tables. Second, nomenclature systems evolve, requiring periodic database updates; we address this through versioned snapshots with timestamps recorded in all logs. Third, symbol harmonization reduces but cannot eliminate all sources of integration bias; it should be combined with other QC steps including batch correction, low-expression filtering, and reference version documentation [19,20]. Finally, our benchmarking focuses on mouse scRNA-seq data processed by CellRanger; the extent of nomenclature drift may differ for other species, sequencing platforms, or processing pipelines, and further validation across these contexts is warranted.

Future work will expand species coverage, provide additional conflict collapsing strategies (sum, max, weighted schemes), develop standardized QC report templates for cross-study comparison, and offer containerized deployment for automated integration pipelines.

## Conclusion

Gene symbol inconsistency causes silent mismatches during RNA-seq data integration, with CellRanger annotation version identified as the primary driver. Across six mouse hippocampus scRNA-seq datasets, harmonization rescued 707–1,098 genes per dataset pair and improved pairwise feature overlap by up to 13.14 percentage points. geneSync provides an offline, auditable solution through hierarchical matching, conflict resolution with full logging, cross-species ortholog mapping, and native single-cell object integration. We recommend incorporating gene symbol harmonization as a routine QC checkpoint prior to large-scale transcriptomic data integration.

## Abbreviations

QC: quality control
NCBI: National Center for Biotechnology Information
scRNA-seq: single-cell RNA sequencing
snRNA-seq: single-nucleus RNA sequencing
GTF: Gene Transfer Format
FASTQ: FASTQ format
HGNC: HUGO Gene Nomenclature Committee
MGI: Mouse Genome Informatics
RGD: Rat Genome Database
ID: identifier
RIKEN: RIKEN (The Institute of Physical and Chemical Research, Japan)
GEO: Gene Expression Omnibus

## Authors’ contributions

ZJ.F. and T.L. conceived and designed the study, developed the geneSync package, and performed the benchmarking analyses. Z.J.F. drafted the manuscript. T.L. reviewed and revised the manuscript.

## Ethics committee approval

Not applicable.

## Acknowledgments

Not applicable.

## Data availability

geneSync is freely available at https://github.com/xiaoqqjun/geneSync under the MIT license. Public benchmarking datasets are accessible through the GEO database via their accession numbers. Due to current data-sharing constraints, the in-house test datasets cannot be released publicly at this stage. The full analysis workflow and scripts are available in the Supplementary Materials.

